# Screening for cryoprotective agent toxicity and toxicity reduction in mixtures at subambient temperatures

**DOI:** 10.1101/2025.05.07.652719

**Authors:** Nima Ahmadkhani, Cameron Sugden, Adam T. Mayo, Adam Z. Higgins

## Abstract

Organ transplantation faces major challenges in preserving and transporting organs due to the limitations of existing cold storage methods. Cryopreservation offers a promising alternative for extending preservation time, but it remains a challenge to avoid toxicity from the high concentrations of cryoprotective agents (CPAs) required to prevent ice formation. In this study, we expanded a previously reported high-throughput CPA toxicity screening platform by retrofitting an automated liquid handling system with subambient cooling capabilities. This enabled systematic assessment of CPA toxicity at 4 °C, a temperature commonly used for CPA equilibration in tissue and organ cryopreservation. Overall, we screened 22 individual CPAs and a wide range of binary mixtures at concentrations up to 12 mol/kg, allowing us to identify CPA combinations that reduce toxicity. Our findings revealed that at 4 °C, CPA toxicity was significantly reduced compared to room temperature. Several CPA combinations resulted in significantly lower toxicity than their constituent CPAs at the same concentration, including 12 CPA mixtures at 6 mol/kg and 8 CPA mixtures at 12 mol/kg. Toxicity neutralization was also observed in 9 cases, especially in combinations involving formamide, acetamide, dimethyl sulfoxide, and glycerol. For example, exposure to 6 mol/kg formamide alone resulted in 20% viability, but the addition of 6 mol/kg glycerol to create a mixture with a total concentration of 12 mol/kg eliminated this toxicity, resulting in a viability of 97%. These findings support the rationale for using multi-CPA cocktails and underscore the potential of rational mixture design to reduce toxicity.

## 1. Introduction

Organ transplantation is a cornerstone of modern medicine, offering life-saving interventions for patients with end-stage organ failure. Since the historic success of the first kidney transplant in 1954, which catalyzed advancements in liver, heart, and pancreas transplants, the field has made remarkable progress [8,22]. However, significant challenges in organ transport and preservation persist.

As of April 2022, over 106,000 patients in the United States were on organ waiting lists, highlighting the urgent need for improved organ availability [8]. Organ transport poses a major challenge, with around 7% of organs encountering issues such as delays and logistical failures between 2014 and 2019, impacting their viability and transplantation success [8]. The viability window for organs like hearts is just 4 to 8 hours, limiting their transport range, while kidneys and pancreases, which can be transported longer distances, often require commercial flights [7,8]. There is a need for enhanced logistical solutions to address these challenges.

Current preservation techniques, including static cold storage and machine perfusion, have limitations. Static cold storage, the conventional method, only allows preservation of hearts for 4 to 8 hours and kidneys for 24 to 36 hours due to ongoing metabolic activity during storage [7,8]. Although machine perfusion can mitigate some preservation-related injuries, it still faces limitations in maintaining long-term organ viability [8]. Consequently, there is a critical need for more advanced preservation techniques to improve transplantation outcomes.

Cryopreservation, which involves cooling biological materials to extremely low temperatures, offers a promising alternative by halting biological processes and enabling long-term storage. This technique is generally divided into slow cooling and vitrification methods. Slow cooling, while effective for suspension-phase cells, poses risks to adherent cells, tissues, and organs due to ice crystal formation. Vitrification, however, employs rapid cooling and high concentrations of CPAs to transition biological materials into a glassy state, thereby avoiding ice formation [11,25]. Vitrification has shown success with diverse biological specimens, from embryos to complex tissues, and has been applied to organ preservation with promising outcomes [9,17,24,27]. For instance, recent studies have reported successful vitrification of rat and rabbit kidneys, maintaining their functionality post-transplantation [14,17,31]. However, challenges persist, particularly with larger human organs, where slower cooling and warming rates necessitate use of higher CPA concentrations to prevent ice formation [15].

Addressing these challenges requires ongoing research into novel CPA formulations that reduce toxicity while enhancing vitrification efficacy. Current research has primarily focused on a limited number of CPAs, which generally share common traits such as low toxicity, cell membrane permeability, and prevention of ice formation [6,18,23,30]. Nonetheless, myriad untested chemicals may offer new opportunities for improving cryopreservation outcomes [1,25,26].

Our previous work evaluated 27 distinct compounds using a fluorescence-based screening approach for rapidly measuring cell membrane permeability and toxicity [1]. The toxicity and permeability characteristics of 23 of the 27 compounds were encouraging. This initial screening approach was limited to solutions containing a single CPA at relatively low concentrations. Therefore, in our recent research, we evaluated the toxicity of 21 of these compounds at higher concentrations and examined the toxicity of binary combinations for 11 compounds. These studies were conducted using an automated liquid handling system to carry out CPA addition and removal at room temperature [2].

In the current study, we modified the automated liquid handling system to allow CPA exposure at subambient temperatures, enabling rapid assessment of CPA toxicity under conditions consistent with those used for CPA equilibration in prior research on tissue and organ cryopreservation [13,17,21]. This experimental setup was used to evaluate the toxicity of 22 compounds at a concentration of 3 mol/kg and temperature of 4 °C. We then selected 14 compounds for toxicity assessment at higher concentrations, both as single CPA solutions and binary CPA mixtures. Our results provide insights into the temperature-dependence of CPA toxicity and highlight the potential for specific combinations of CPAs to reduce toxicity.

## 2. Materials and methods

### 2.1. Materials

We used 22 CPAs in this study (see Table 1). These CPAs were selected from the 27 CPAs previously screened by Ahmadkhani et al. [1] due to their favorable permeability and toxicity. Additional chemicals and reagents—including PrestoBlue, HEPES-buffered saline (HBS), Dulbecco’s Modified Eagle Medium (DMEM), fetal bovine serum, and penicillin-streptomycin— were procured and prepared as described by Warner et al. [30].

**Table 1.** CPA exposure conditions tested at 4 ± 2°C.

| CPAs tested at 3 mol/kg (10 and 30 min exposure) | CPAs tested at 6 and 12 mol/kg (30 min exposure) |
| --- | --- |
| 1,3-Dihydroxyacetone (DHA) | 1,3-Propanediol (PD) |
| 1,3-Propanediol (PD) | 2,3-Butanediol (BD) |
| 2,3-Butanediol (BD) | 2-Methoxyethanol (ME) |
| 2-Methoxyethanol (ME) | 2-Methyl-1,3-Propanediol (MP) |
| 2-Methyl-1,3-Propanediol (MP) | Acetamide (AM) |
| 2-Methyl-2,4-Pentanediol (MPD) | Diethylene Glycol (DG) |
| Acetamide (AM) | Dihydroxyacetone (DHA) |
| Diethylene Glycol (DG) | Dimethyl Sulfoxide (DMSO) |
| Diglyme | Dimethylacetamide (DMA) |
| Dimethyl Sulfoxide (DMSO) | Ethylene Glycol (EG) |
| Dimethylacetamide (DMA) | Formamide (FA) |
| Ethylene Glycol (EG) | Glycerol (Gly) |
| Formamide (FA) | N-Methylacetamide (NMA) |
| Glycerol (Gly) | Propylene Glycol (PG) |
| N-Methylacetamide (NMA) | Binary CPA mixtures composed of the 14 CPAs above (equimolar split between CPAs). We tested 87 binary mixtures at 6 mol/kg and 82 binary mixtures at 12 mol/kg. |
| Propionamide (PA) |  |
| Propylene Glycol (PG) |  |
| Tetraethylene Glycol Dimethyl Ether (TGDE) |  |
| Tetrahydrofurfuryl Alcohol (TFA) |  |
| Triethylene Glycol (TG) |  |
| Triethylene Glycol Diacetate (TGD) |  |
| Triglyme |  |

### 2.2. Experimental Overview

This study builds on our prior work, which assessed CPA toxicity at room temperature [2,30]. As in our previous studies, we used automated liquid handling for CPA addition and removal, enabling precise and reproducible solution exchanges. To maintain subambient temperatures, we used a custom-built plate cooling module designed to fit within the deck layout of the Hamilton Microlab STARlet, with designated positions for both deep-well plates and assay plates (MéCour Temperature Control, LLC). The plate cooler was connected to a circulating bath, which maintained the system at the desired temperature of 4 °C throughout the liquid handling process.

This experimental platform was employed to evaluate 22 CPAs and their binary mixtures at various concentrations (219 compositions in total).

Bovine pulmonary artery endothelial cells (BPAEC) were cultured in 96-well plates and used to evaluate CPA toxicity, following the same general protocol as in our prior research [30]. Cell viability was first measured before CPA treatment using the PrestoBlue assay, which relies on the metabolic conversion of resazurin into resorufin by living cells. After this baseline measurement, CPAs were introduced using the automated liquid handling platform. A second PrestoBlue viability reading was taken 20-24 hours post-exposure. To quantify toxicity, we applied a dual normalization method: each post-treatment PrestoBlue reading was normalized to the pre-treatment value from the same well, as well as the average signal from the untreated control wells. The well plate layout was modified slightly compared to our previous study [30] to increase the number of compositions that could be tested per plate. We tested 19 CPA compositions per plate (4 wells each), with the remaining wells used for controls: CPA-free positive control (8 wells), negative control (4 wells), and cell-free background control (8 wells).

In order to improve efficiency, the CPA addition and removal procedures were streamlined compared to those used in our previous study [30]. Given that EG was identified as a low-toxicity CPA in both our results and prior research [20,28,30], we opted to use 3 mol/kg EG for intermediate steps before and after introducing a higher CPA concentration to the cells. Methodological details for CPA addition and removal are provided in the Supplementary Material, along with predictions of cell volume changes (Figure S.1) and experimental results comparing different CPA addition and removal methods (Figure S.2-S.3).

To prepare CPA solutions, isotonic HBS was used to prepare 18 mol/kg stock solutions of the 22 CPAs, and these stock solutions were combined volumetrically with HBS to achieve the desired CPA concentration. Due to an oversight, the HBS used for these experiments was not pH-adjusted to 7.3. Subsequent testing, using identically prepared buffer, revealed the actual pH to be approximately 5.5. This deviation did not appear to negatively affect cell health: viability outcomes in this study were consistent with those from our prior study that used pH-adjusted buffers [30]. Specifically, the isotonic control wells showed robust cell growth, as evidenced by a 2-fold increase in the PrestoBlue signal after 24 hours, suggesting that the cells roughly doubled in number despite short-term exposure to lower pH conditions.

### 2.3. Statistical analysis

Statistical analyses were performed using the Python packages statsmodels and SciPy. Consistent with our previous study [30], outliers were identified using the boxplot method based on the standard interquartile range (IQR) criterion, where values falling outside 1.5 times the IQR from the first and third quartiles were considered outliers. Data are presented as mean ± standard error of the mean (SEM). In most cases, one-way analysis of variance (ANOVA) was used for comparisons among multiple groups, followed by Tukey’s post hoc test for pairwise comparisons. A p-value below 0.05 was considered statistically significant. We used a different approach to analyze toxicity reduction in CPA mixtures, due to the large number of comparisons. In this case, Welch’s two-sample, two-sided t-test was performed, followed by the Benjamini–Hochberg method as a correction for multiple testing [5]. A corrected p-value (q-value) < 0.05 was considered statistically significant. The normality assumption for the data was assessed using Q-Q plots. Visual inspection of the Q-Q plots indicated that the normality assumption was valid for all groups analyzed.

## 3. Results

To enable high-throughput screening of CPA mixtures, we automated CPA exposure and multistep CPA addition and removal using a Hamilton Microlab STARlet liquid handling system (Figure 1). To maintain subambient conditions, we modified our existing system [30] by adding a plate cooling module with a circulating bath set to –4 °C, which stabilized the plate temperature at 4 ± 2 °C. CPA solutions were dispensed into deep-well plates according to a randomized 96-well format and subsequently transferred to assay plates containing bovine pulmonary artery endothelial cells to assess CPA toxicity. Viability was measured 20–24 hours post-exposure using the PrestoBlue assay.

**Figure 1.**
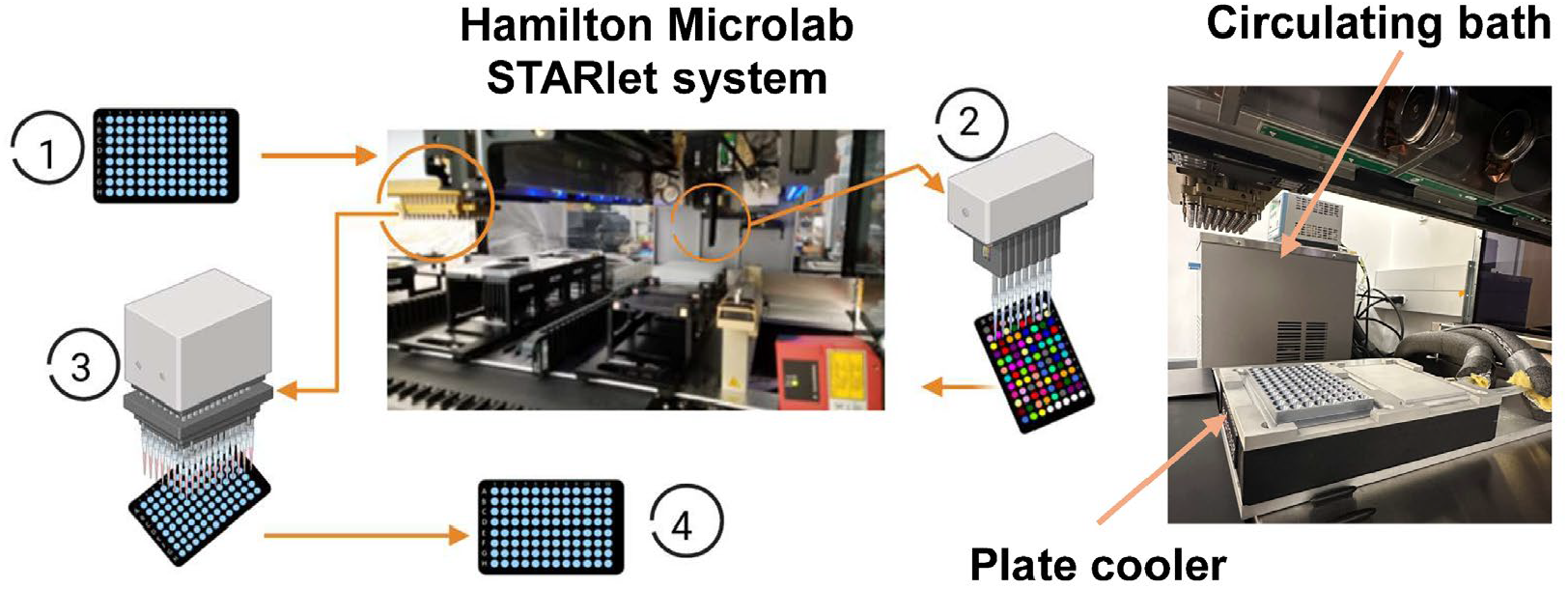
Automated liquid handling setup with temperature control. The experimental workflow consists of: (1) assess cell viability prior to CPA exposure using the PrestoBlue assay; (2) dispense CPA solutions into randomized locations in a deep well plate using 8 independent pipettors; (3) transfer CPAs from the deep well plate to the assay plate containing cells using the 96 channel head; (4) assess cell viability 20–24 hours after CPA exposure using the PrestoBlue assay. Temperature was maintained at 4 °C using a plate cooling module with a circulating bath. This figure was created in Biorender.

### 3.1. Toxicity of CPA solutions at 4 °C

We began by assessing the toxicity of the 22 CPAs that we found to have high cell membrane permeability and low toxicity at 0.7 mol/kg in our previous study [1]. These CPAs were tested at a concentration of 3 mol/kg for two exposure times: 10 min and 30 min (Figure 2). Several CPAs, such as diglyme, triglyme, MPD, TGDE, TGD, TFA, and PA, exhibited significant toxicity over both exposure periods. However, there were numerous CPAs that resulted in high cell viability. In general, cell viability after a 10 min exposure was comparable to or greater than the cell viability after a 30 min exposure, indicating greater toxicity for longer exposure times. Based on the results in Figure 2, we selected 14 CPAs for subsequent testing at higher concentrations.

**Figure 2.**
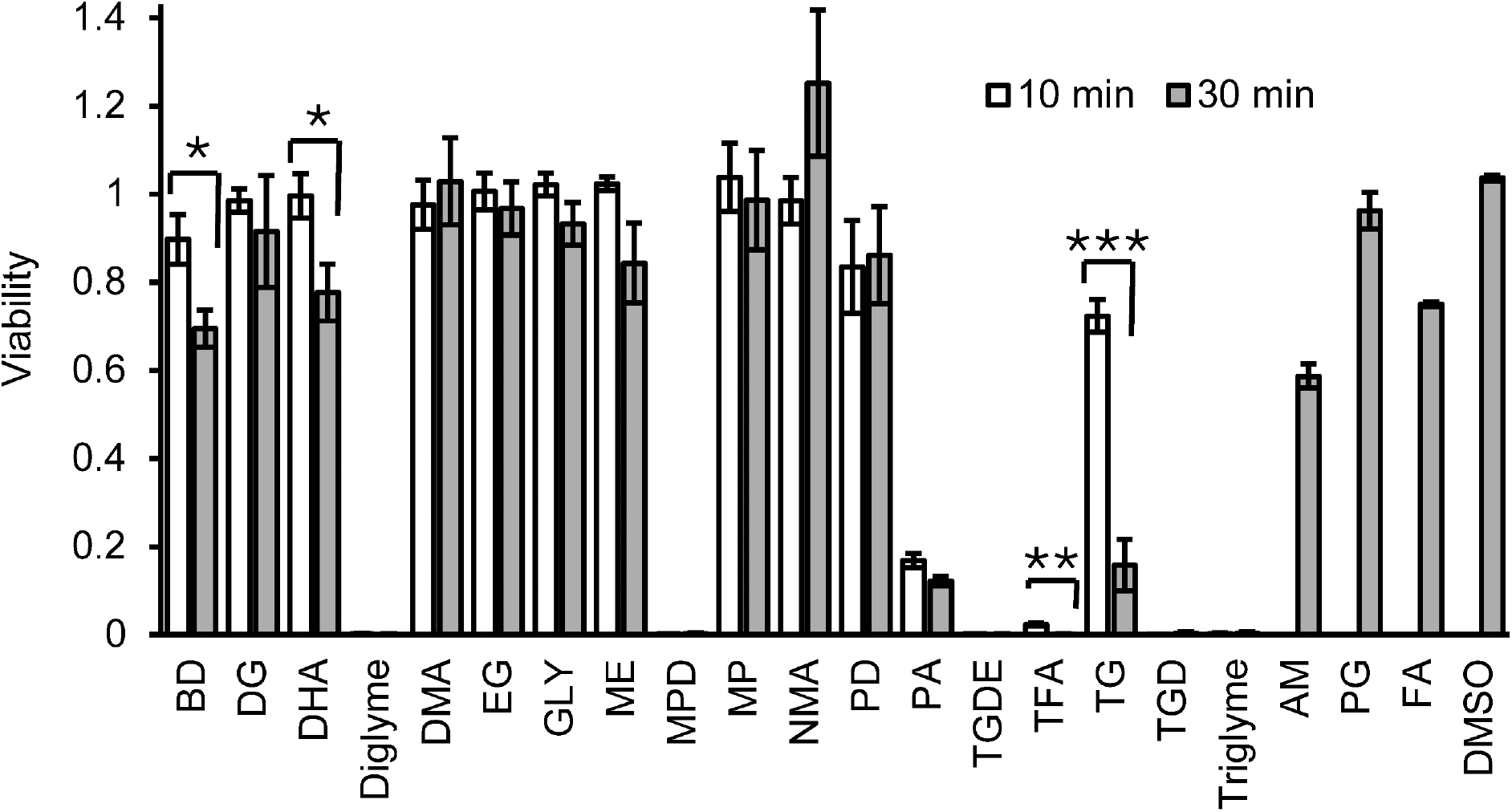
Effect of exposure time on cell viability after exposure to 3 mol/kg CPA. Each bar represents the mean ± SEM from 3–7 replicate wells. Statistically significant differences between groups are indicated by asterisks, with the following p-value thresholds: * p < 0.05, ** p < 0.01, *** p < 0.001. No data was collected for AM, PG, FA and DMSO at the 10 min exposure time.

For CPA concentrations of 6 and 12 mol/kg, we tested 30-min exposure to 14 individual CPAs as well as their binary mixtures, resulting in around 100 compositions at each concentration. The results of these experiments are shown in Figures 3 and 4. At a concentration of 6 mol/kg, 42% of the CPA compositions yielded a cell viability exceeding 80%. Conversely, exposure to 12 mol/kg CPA resulted in greater levels of toxicity, indicating that cytotoxicity increases in a concentration-dependent manner.

**Figure 3.**
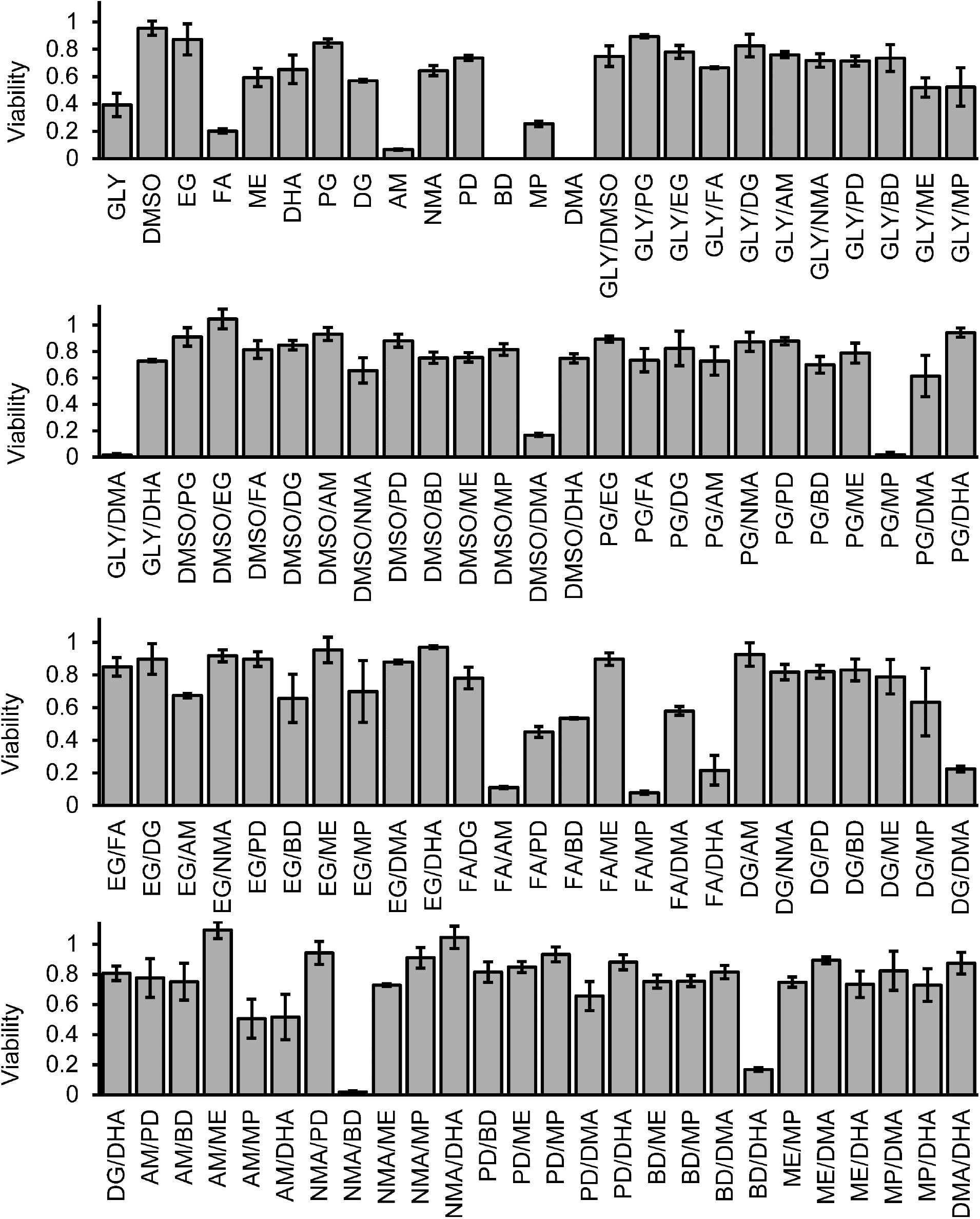
Cell viability after exposure to 6 mol/kg CPA mixtures for 30 min. Each bar represents the mean ± SEM from 3–4 replicate wells.

**Figure 4.**
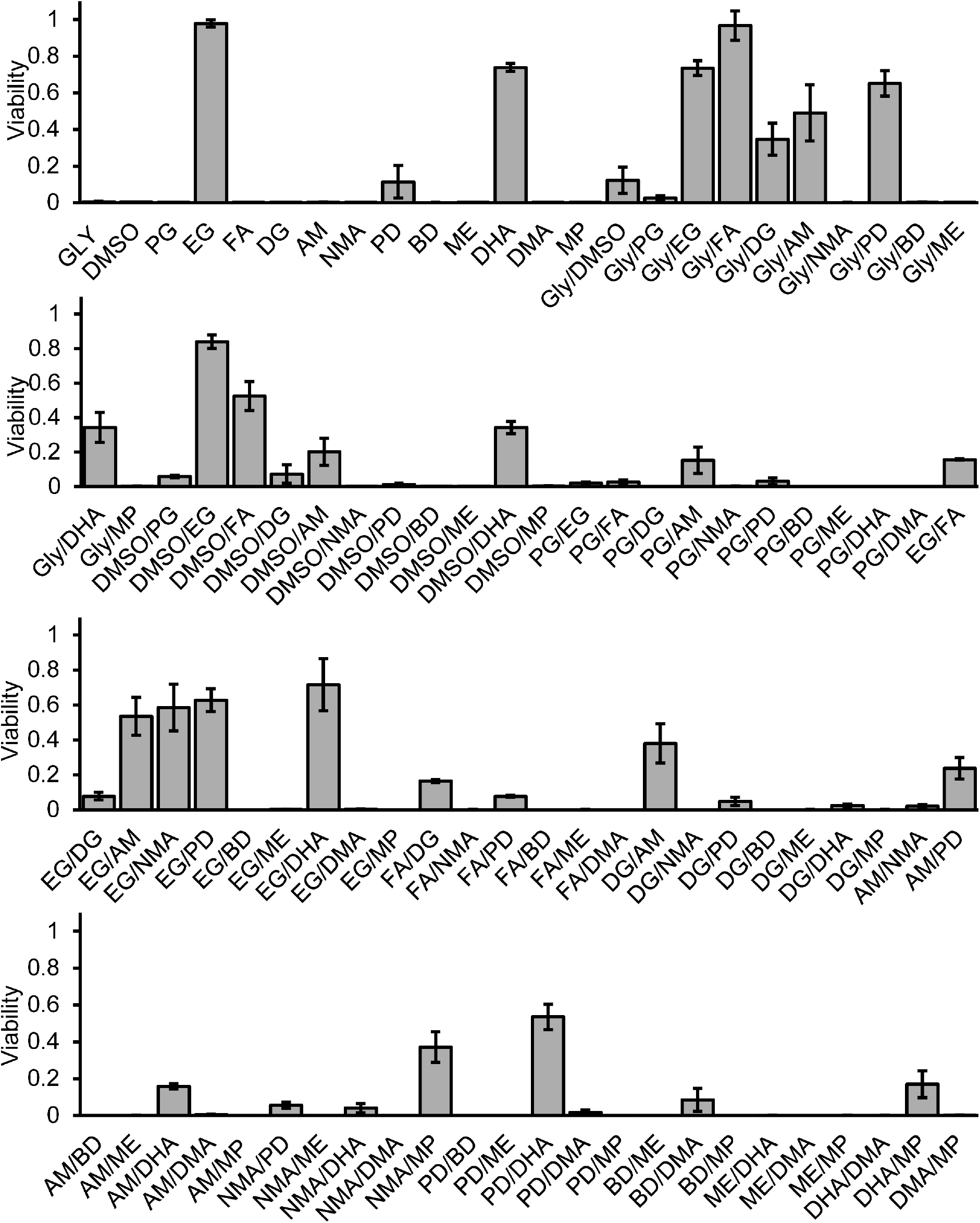
Cell viability after exposure to 12 mol/kg CPA mixtures for 30 min. Each bar represents the mean ± SEM from 3–12 replicate wells.

### 3.2. Temperature dependence of CPA toxicity

To assess the effects of temperature on CPA toxicity, we compared our results at 4 °C to the results of our previous study, which assessed toxicity at room temperature [2]. Figure 5 shows cell viability results for 54 CPA compositions at both 4 °C and room temperature. In both cases the CPA solutions were tested at a concentration of 6 mol/kg. The results demonstrate a substantial disparity in the survival rate depending on temperature. For example, PG showed full toxicity at room temperature, but only minor toxicity at a temperature of 4 °C. Generally, the ability of the cells to survive CPA exposure was greater at 4 °C than at ambient temperature. Statistical analysis indicated that 80% of the compounds exhibited significantly greater viability at 4 °C compared to room temperature.

**Figure 5.**
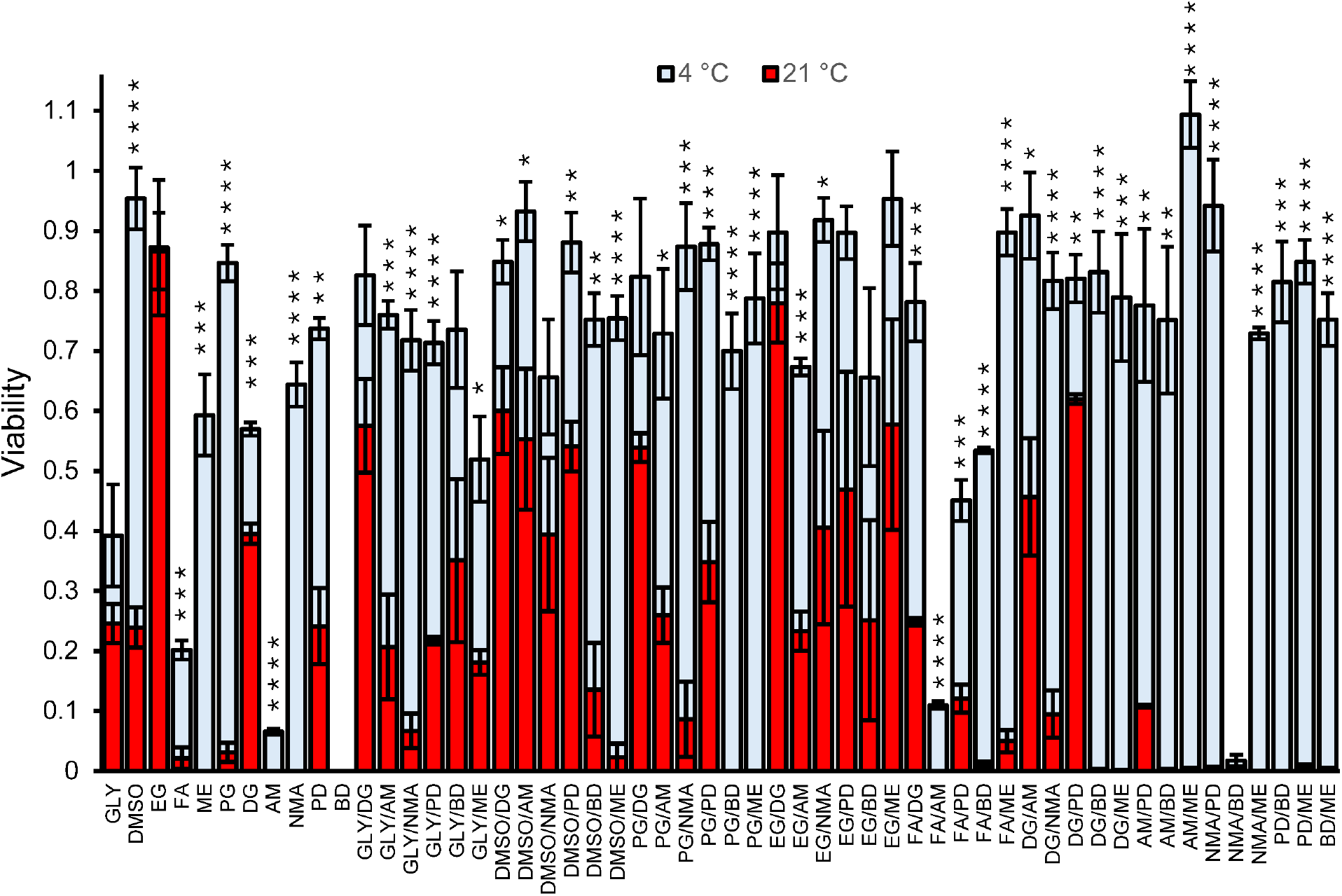
Temperature dependence of CPA toxicity for 30 min exposure to 6 mol/kg CPA. Bars represent the mean ± SEM from 3–4 replicate wells. Results at room temperature (∼21 °C) are from our previous study [2]. Statistically significant differences are indicated by asterisks: * p < 0.05, ** p < 0.01, *** p < 0.001, **** p < 0.0001.

### 3.3. Toxicity reduction in CPA mixtures

Prior studies suggest that CPA cocktails may have a reduced toxicity compared to individual CPAs [2,3,4,10,12,19,30]. To probe for possible toxicity reduction in mixtures, we compared binary CPA mixtures to their corresponding single-CPA solutions at the same total CPA concentration. Notably, 47 of the 87 mixtures tested at 6 mol/kg resulted in higher cell viability than both corresponding single-CPA solutions. In contrast, only 2 mixtures exhibited lower viability than both single CPA solutions. Together, this suggests a general trend of toxicity reduction in CPA mixtures. As shown in Figure 6, 12 mixtures exhibited statistically significant toxicity reduction characterized by significantly higher cell viability for the mixture than both single CPA solutions.

**Figure 6.**
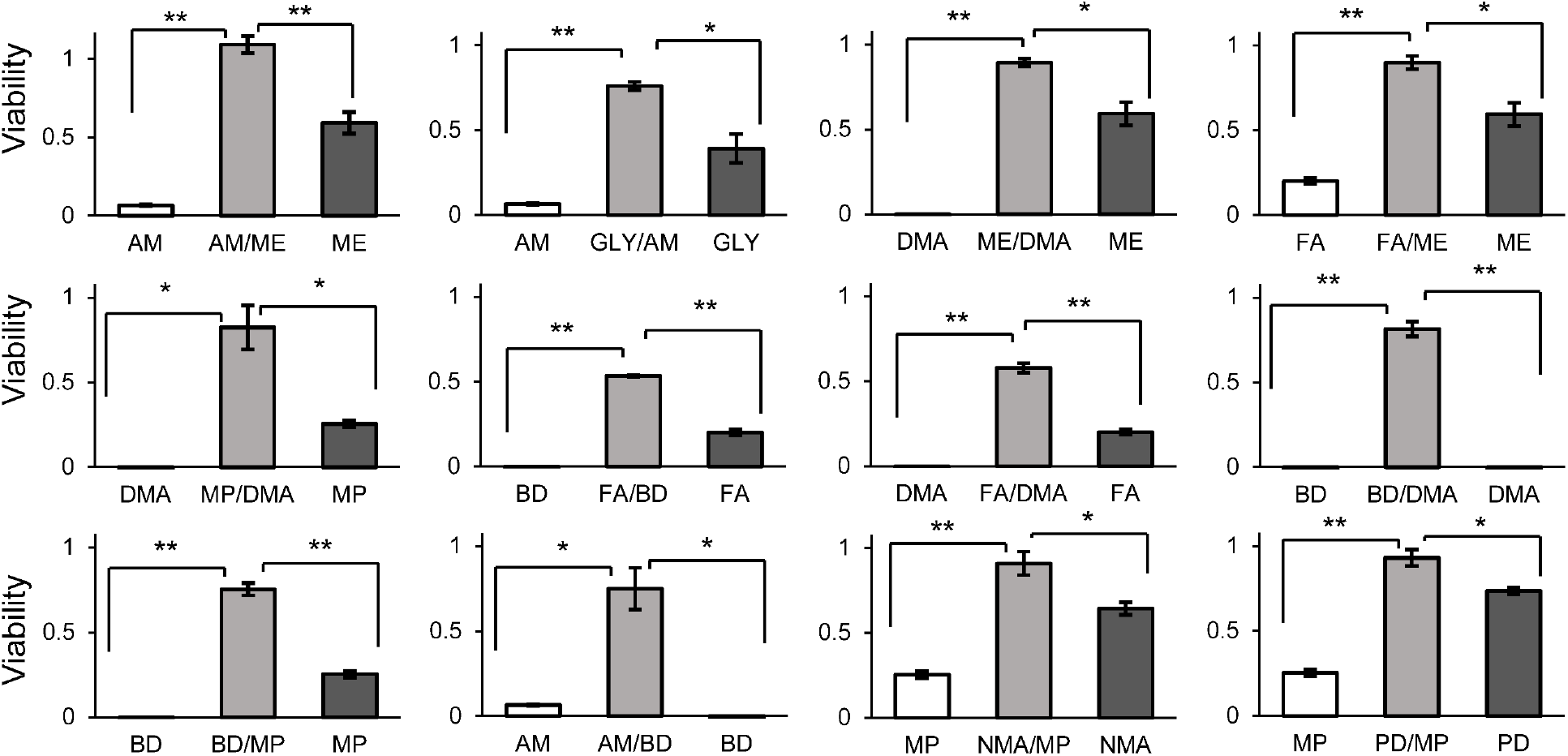
Toxicity reduction in 6 mol/kg CPA mixtures. To identify CPA combinations that reduce toxicity, we compared each mixture to its 6 mol/kg individual CPA constituents. Each bar represents the mean ± SEM from 3–4 replicate wells. Statistically significant differences between groups are indicated by asterisks, with the following p-value thresholds: * p < 0.05, ** p < 0.01.

We also explored toxicity reduction for the 12 mol/kg CPA mixtures. At this concentration, we tested 82 binary combinations. To assess toxicity reduction, we limited our analysis to experimental groups in which at least one composition (either the CPA mixture or one of the individual CPAs) resulted in a cell viability greater than 2%. A total of 50 mixtures met this criterion. Among these, 17 CPA mixtures showed higher viability than both of the corresponding single CPA solutions, indicating possible protective effects for 34% of the CPA mixtures. In contrast, there were no cases where the viability of the mixture was lower than both single CPA solutions. As shown in Figure 7, statistical analysis revealed 8 cases where toxicity was significantly lower for the mixture. In some cases, such as GLY/FA, cell viability was high for the mixture, but both single CPA solutions were completely toxic.

**Figure 7.**
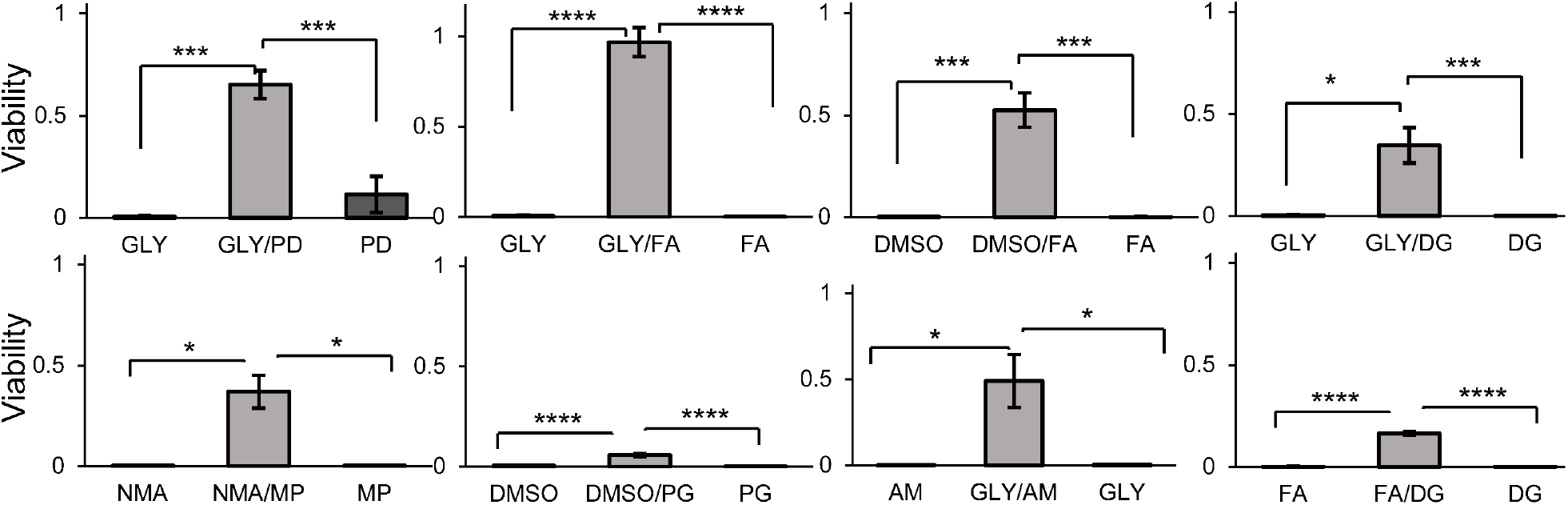
Toxicity reduction in 12 mol/kg CPA mixtures. To identify CPA combinations that reduce toxicity, we compared each mixture to its 12 mol/kg individual CPA constituents. Each bar represents the mean ± SEM from 3–10 replicate wells. Statistically significant differences between groups are indicated by asterisks, with the following p-value thresholds: * p < 0.05, ** p < 0.01, *** p < 0.001, **** p < 0.0001.

### 3.4. Toxicity neutralization

To delve deeper into the toxicity of multi-CPA mixtures, we analyzed toxicity neutralization as described by Fahy [10]. Fahy showed that formamide toxicity can be neutralized by adding DMSO, resulting in a DMSO/FA mixture with a higher total concentration but lower toxicity. We explored toxicity neutralization for both 6 mol/kg and 12 mol/kg mixtures. The 6 mol/kg mixtures were comprised of 3 mol/kg concentrations of each CPA. Thus, to probe for toxicity neutralization in the 6 mol/kg mixtures, we compared the 6 mol/kg mixture to the individual CPAs at a concentration of 3 mol/kg. Similarly, for analysis of the 12 mol/kg mixtures, we compared the mixture to the individual CPAs at a concentration of 6 mol/kg. In both cases, a higher viability for the mixture indicates toxicity neutralization. We observed statistically significant toxicity neutralization in 18 cases, but for 9 of these cases the viability for the mixture was very low (<2%). The remaining 9 cases are depicted in Figures 8 and 9. The most prominent effect was obtained when using a combination of FA and GLY at a concentration of 12 mol/kg. In this case, the 12 mol/kg mixture resulted in 97% viability, which is much higher than the 20% viability observed for 6 mol/kg FA alone. This indicates that GLY neutralizes FA toxicity. DMSO and FA also exhibited toxicity neutralization, consistent with Fahy’s previous observations [10]. Overall, the results highlight possible toxicity neutralizing effects for several CPA combinations.

**Figure 8.**
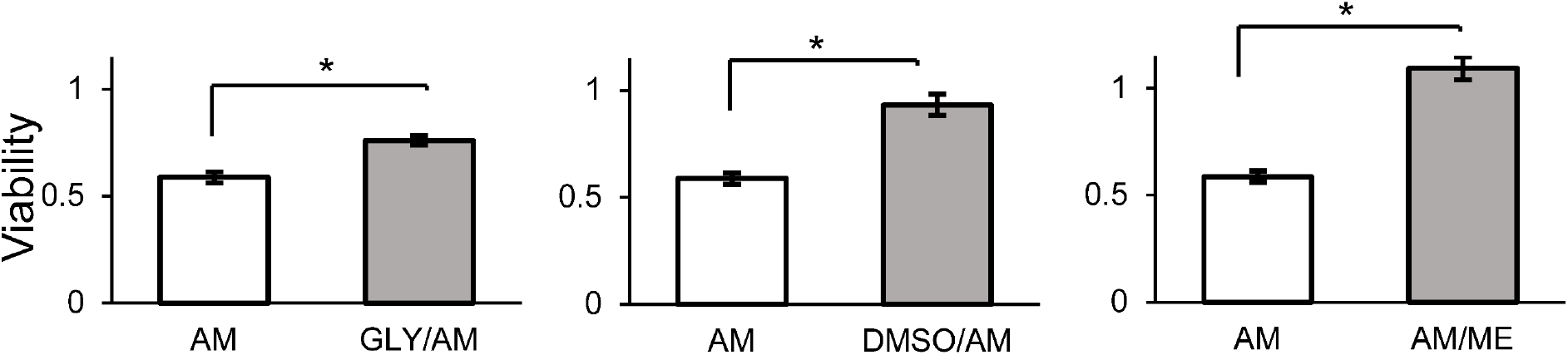
Toxicity neutralization in 6 mol/kg CPA mixtures. If the 6 mol/kg mixture has higher viability than one of the single CPAs at a 3 mol/kg concentration, this indicates toxicity neutralization. Each bar represents the mean ± SEM from 3–4 replicate wells. Statistically significant differences between groups are indicated by an asterisk: * p < 0.05.

**Figure 9.**
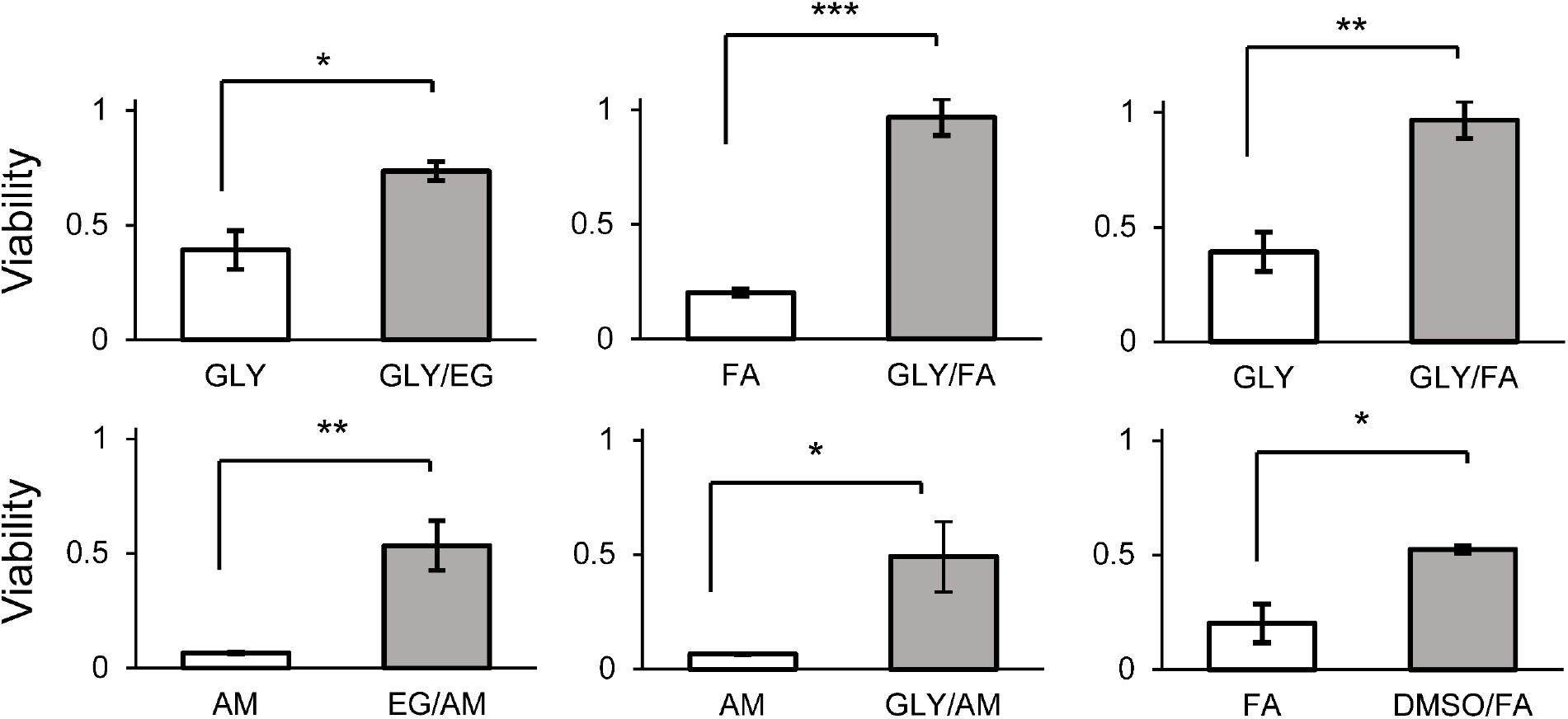
Toxicity neutralization in 12 mol/kg CPA mixtures. If the 12 mol/kg mixture has higher viability than one of the single CPAs at a 6 mol/kg concentration, this indicates toxicity neutralization. Each bar represents the mean ± SEM from 3–8 replicate wells. Statistically significant differences between groups are indicated by asterisks, with the following p-value thresholds: * p < 0.05, ** p < 0.01, *** p < 0.001.

## 4. Discussion

In this study, we expanded our previously reported high-throughput CPA toxicity screen [30] by retrofitting the automated liquid handler with a plate cooling module for subambient temperature control. Unlike our earlier works—where automated screening was performed at room temperature [2,30]—the current study examined CPA toxicity at 4 °C, the temperature most commonly used for CPA equilibration in organ and tissue cryopreservation [13,17]. This allowed us to evaluate CPA toxicity under more relevant, subambient conditions and revealed a pronounced temperature dependence of toxicity. For example, several CPA compositions yielded > 80% viability at 4 °C but were completely toxic at room temperature.

Moreover, by testing 22 individual CPAs at 3 mol/kg and 14 CPAs and their binary mixtures at 6 and 12 mol/kg, we probed toxicity reduction in mixtures in greater breadth than previous studies [4,10]. Toxicity reduction in CPA mixtures is thought to occur through two primary mechanisms: mutual dilution and toxicity neutralization. In mutual dilution, each CPA in the mixture decreases the effective concentration of the others. Since higher CPA concentrations are typically more toxic, this dilution effect can make the overall mixture less harmful than the same concentration of each CPA on its own [11,29,30]. In contrast, toxicity neutralization refers to a more specific interaction where the presence of one CPA directly counteracts or eliminates the toxic effects of another [11].

In this study, we first examined toxicity reduction by comparing binary CPA mixtures to their constituent CPAs at the same total concentration. This approach does not distinguish between mutual dilution and toxicity neutralization as the mechanism of toxicity reduction. We observed higher viability in 54% of mixtures at 6 mol/kg and in 34% of mixtures at 12 mol/kg, suggesting possible widespread toxicity reduction in mixtures. Statistical analysis revealed that toxicity was significantly reduced in 14% of mixtures at 6 mol/kg and 16% of mixtures at 12 mol/kg. These results reinforce the common practice of using multi-CPA cocktails (e.g., VS55, DP6, M22) to reduce toxicity [13,16,17].

To examine toxicity neutralization, we compared binary CPA mixtures to their constituent CPAs at half the mixture’s concentration. Toxicity neutralization was particularly notable for GLY and FA, which is consistent with our previous report [2]. In particular, we observed that the addition of 6 mol/kg GLY to 6 mol/kg FA increased viability from 20% to 97%, despite the overall higher CPA concentration in the GLY/FA mixture. This indicates that GLY neutralizes FA toxicity. In addition, we found that DMSO neutralized the toxicity of FA and AM, which aligns with Fahy’s findings in renal cortical slices [10]. Overall, the amides FA and AM demonstrated neutralization with four partners (GLY, DMSO, ME, EG), suggesting that incorporating specific amides into vitrification cocktails could be a powerful strategy to balance CPA permeability, glass-forming ability, and cytotoxicity.

Looking forward, our results pave the way for targeted mixture design and higher-order screening to identify minimal-toxicity vitrification cocktails. This will require assessment of the concentration needed for vitrification for each CPA combination, so that low toxicity compositions can be found at this threshold concentration. In this study, the highest CPA concentration we tested was 12 mol/kg, which corresponds to concentrations ranging from 37% w/v to 63% w/v, depending on the CPA. This exceeds the threshold concentration for vitrification for several individual CPAs, including DMSO and PG [11]. For most of the CPA mixtures, the threshold concentration is not available in literature. This highlights the need for future studies to assess the concentration needed to vitrify in CPA mixtures. Ultimately, by screening CPAs at the temperatures and concentrations used in practice, we can more reliably predict and mitigate cryoinjury, accelerating the development of robust, clinically relevant cryopreservation solutions.

## 5. Conclusions

In this study, we expanded our previous toxicity screening method to include subambient temperature control, enabling systematic assessment of the toxicity of 22 individual CPAs and a broad array of binary mixtures at 4 °C. Our results demonstrate a pronounced temperature dependence of CPA toxicity, as evidenced by significantly higher viability at 4 °C than room temperature for 43 of the 54 CPA compositions tested. This supports the established practice of performing CPA equilibration at subambient temperatures and underscores the value of conducting toxicity screening under these conditions to identify low-toxicity formulations. Our results also provide strong evidence for widespread protective effects in CPA mixtures that reduce overall mixture toxicity. We quantified this by comparing binary CPA mixtures to both individual CPAs at the same total concentration. Higher viability was observed in 54% of mixtures at 6 mol/kg and 34% of mixtures at 12 mol/kg, and there was a total of 20 cases where the toxicity reduction in the mixture was statistically significant. We also observed toxicity neutralization in several cases, defined as lower toxicity in the mixture than one of the individual CPAs at half the total mixture concentration. Notably, the toxicity of amides such as formamide and acetamide was mitigated not only by DMSO, but also by other CPAs, including glycerol. To further advance from these encouraging findings, future investigations should focus on augmenting the chemical repository of CPAs by including a wider array of compounds that may exhibit low toxicity or protective properties. Moreover, the vast amount of data produced by high-throughput screening can be used to create predictive models that would enhance the efficiency of identifying and assessing novel CPAs. These developments are anticipated to enhance vitrification methods, resulting in improved preservation of intricate biological structures such as organs and wider applicability in medical and scientific domains. Further advancements in this field have the potential to greatly improve the effectiveness of organ transplantation.

## Supporting information

Viability Data

## 6. Acknowledgements

This work was supported by a generous donation to the laboratory of AZH from the Cryonics Institute.

## 8. Declaration of generative AI and AI-assisted technologies in the writing process

During the preparation of this work, the authors used ChatGPT to assist with wording and sentence structure. After using this tool/service, the authors reviewed and edited the content as needed and take full responsibility for the content of the publication.

