## Supplementary Material for "Screening for cryoprotective agent toxicity and toxicity reduction in mixtures at subambient temperatures"

Nima Ahmadkhani[1], Cameron Sugden[1], Adam T. Mayo[1], Adam Z. Higgins[1]
[1]School of Chemical, Biological and Environmental Engineering, Oregon State University.

**Addition and removal of 6 mol/kg CPA**

Figure S.1 shows normalized cell-volume trajectories for several different CPA addition and removal protocols predicted using a biophysical model of cell membrane transport [1]. Figure S.2 shows the corresponding cell viability that we measured for each of these methods. For the multistep methods, CPA addition and the first step of CPA removal were carried out at 4 °C. The remaining CPA removal steps were carried out at room temperature (~22 °C) to reduce the potential for osmotic damage. We considered two approaches, the standard multi-step approach and an approach we refer to as the EG method. In the standard approach, the CPA concentration is incrementally increased during CPA addition and incrementally decreased during CPA removal. The EG method uses a similar multi-step approach, but EG is used for all intermediate steps. The EG method is advantageous because EG is relatively nontoxic, and because this method uses the same intermediate steps for all CPAs, simplifying the experimental workflow. We also considered "short" and "long" methods, in which the durations of the intermediate steps were varied. Figure S.1 shows volume predictions for glycerol and formamide, the CPAs with the lowest and highest cell membrane permeability values from our previous study [1]. For glycerol, volume predictions suggest possible excessive cell shrinkage during the first step of CPA addition using the standard method, and possible excessive swelling during the first step of CPA removal using the short EG method. This is due to glycerol's low permeability. For formamide, none of the methods are expected to produce cell volume changes that exceed the osmotic tolerance limits. However, it should be noted that the EG method is predicted to cause cell swelling upon exposure to 6 mol/kg formamide. This is counterintuitive because the 6 mol/kg formamide solution is relatively hypertonic compared to the 3 mol/kg EG from the preceding step. The swelling is caused by the relatively rapid permeation of FA into the cell, and the relatively slow egress of intracellular EG. This highlights the possibility of even greater swelling for CPAs that permeate faster than FA.

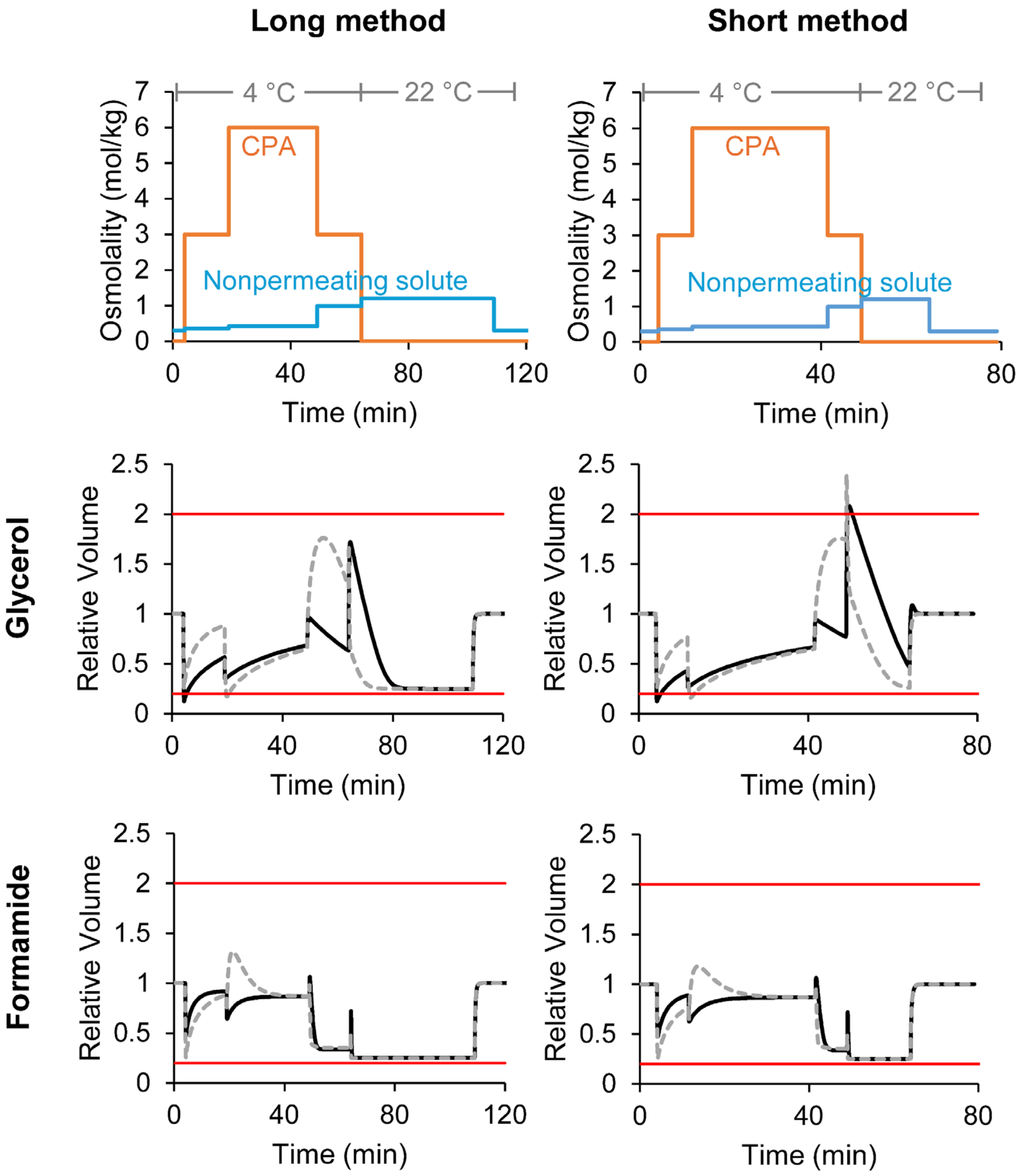


*Figure S.1. Simulated cell-volume dynamics using a MATLAB-based biophysical model [1]. The top panels show the temperature and composition during CPA addition and removal using the long and short methods. The middle panels show predictions of osmotically active cell volume for addition and removal of 6 mol/kg glycerol. The bottom panels show predictions of osmotically active cell volume for addition and removal of 6 mol/kg formamide. The black lines show "standard" CPA addition and removal, the gray dashed lines show CPA addition and removal using the "EG method", and the red lines show the osmotic tolerance limits [2].*

As shown in Figure S.2, multistep CPA addition and removal protocols led to substantially higher cell viability than the single-step method across all tested CPAs at 6 mol/kg. This underscores the need for multi-step methods to prevent osmotic damage. The short multi-step methods consistently outperformed or matched the viability outcomes of their long counterparts, regardless of CPA type. Notably, the short EG protocol yielded comparable viability to the standard short method, with the exception of ME. In this case, the viability for the standard short method was significantly higher than the viability for the short EG method ($p < 0.05$). These findings underscore the efficiency of short, multistep protocols in mitigating osmotic stress and enhancing cell survival. Because the EG method obviates the need for custom intermediate solutions, we adopted the short EG protocol for all subsequent experiments.

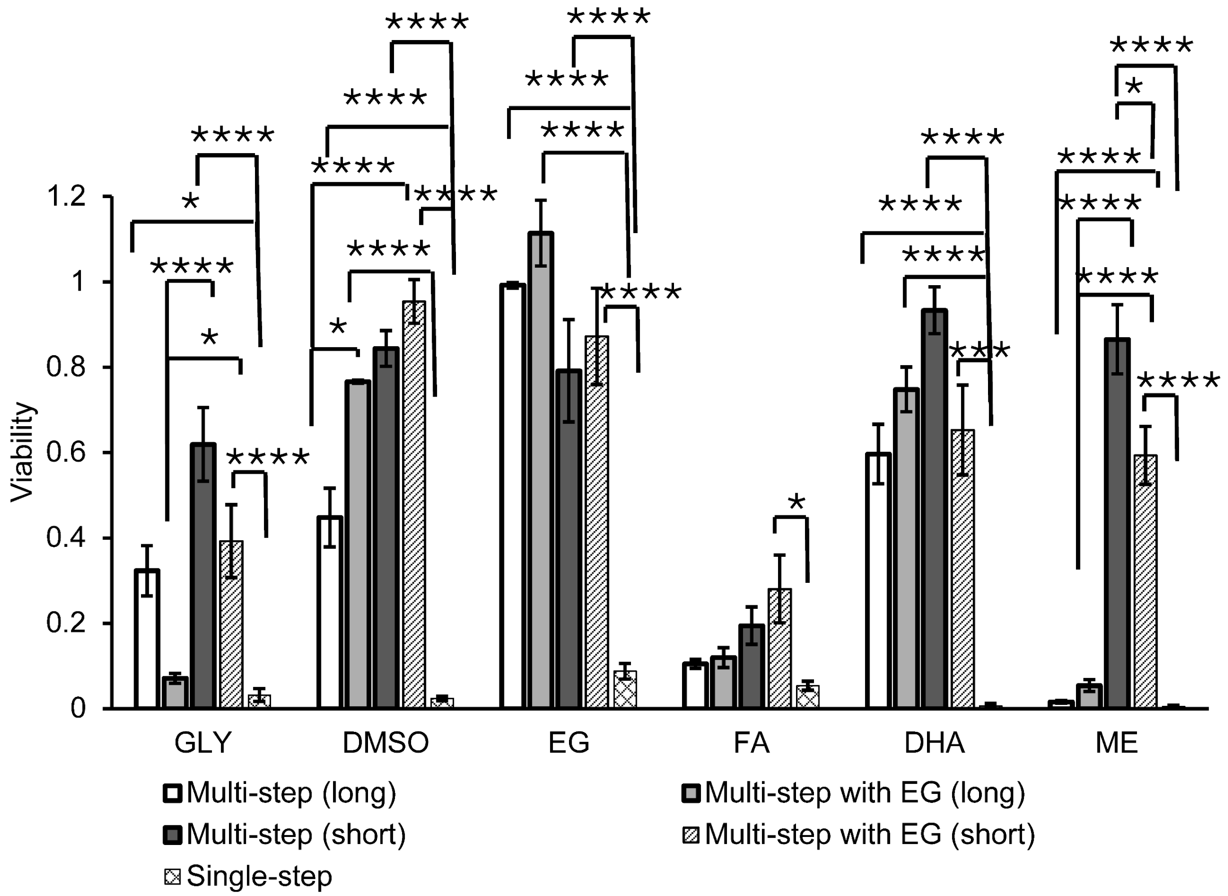


*Figure S.2. Comparison of CPA addition and removal methods for exposure to 6 mol/kg CPA. Each bar represents the mean ± SEM from 3–4 replicate wells. Statistically significant differences between groups are indicated by asterisks, with the following p-value thresholds: * p < 0.05, ** p < 0.01, *** p < 0.001, ****p< 0.0001.*

**Addition and removal of 12 mol/kg CPA**

Figure S.3 shows cell viability results after addition and removal of 12 mol/kg CPA using the methods depicted in Figure S.4. Among the five CPAs shown at Figure S.3, three resulted in near-zero viability across all methods, indicating complete toxicity under the conditions used. For the remaining CPAs (i.e., DHA and EG), significant differences were observed between protocols. In particular, the short method with EG consistently produced the highest cell viability. Comparisons also showed that using EG for intermediate steps improved viability compared to standard CPA addition and removal for DHA, and the short protocol outperformed the long protocol for both DHA and EG. These results highlight the importance of optimizing both exposure duration and solution composition to minimize osmotic damage and toxicity during CPA addition and removal. Based on these results, we chose to use the short EG method for all subsequent experiments.

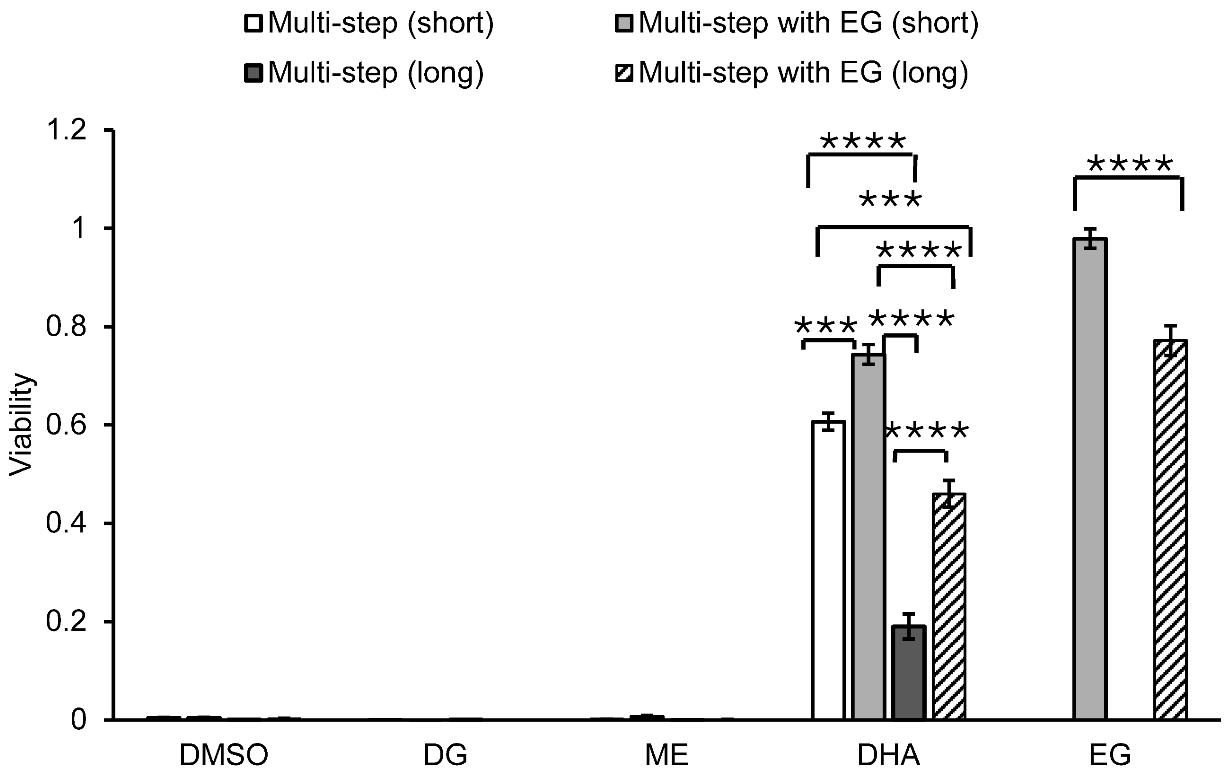


*Figure S.3. Comparison of CPA addition and removal methods for exposure to 12 mol/kg CPA. Each bar represents the mean ± SEM from 3–4 replicate wells. Experiments were conducted at 4 °C. Statistically significant differences between groups are indicated by asterisks, with the following p-value thresholds: * $p < 0.05$, ** $p < 0.01$, *** $p < 0.001$, ****$p < 0.0001$.*

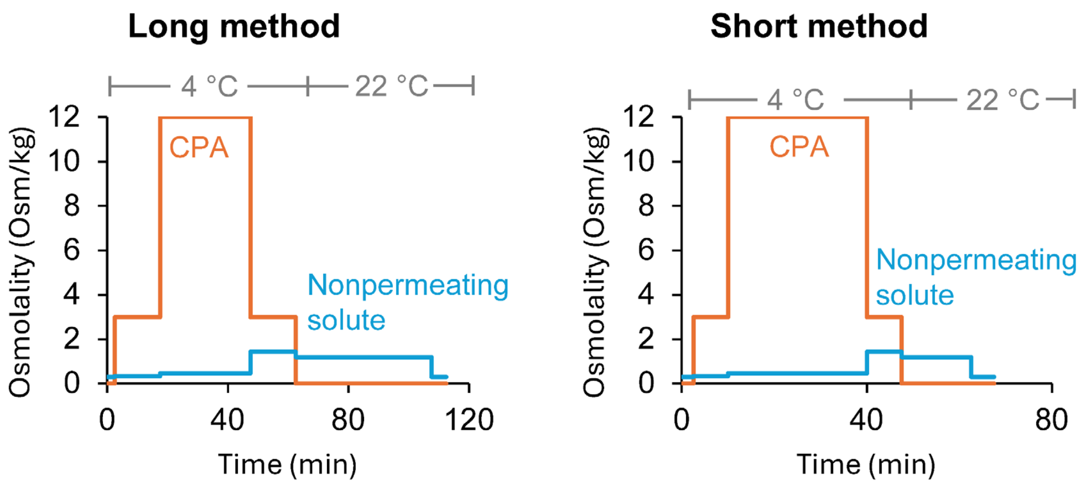


*Figure S.4. Protocols for addition and removal of 12 mol/kg CPA, comparing short and long methods. The orange lines represent the concentration profile of permeating CPAs over time, while the blue lines indicate non-permeating solute concentrations.*
